# Benchmarking Universal Single-Copy Orthologs for Phylogenomics: The Impact of Paralogy

**DOI:** 10.64898/2026.09.25.754399

**Authors:** Miguel Aranda, Nikita Kulikov, Iker Irisarri

## Abstract

Universal Single-Copy Orthologs (USCOs) are taxonomically-defined gene sets originally proposed to evaluate completeness in genome assemblies. Due to their single- or low-copy nature, they have also been proposed as ideal markers for phylogenomics. Here, we test this assumption using two vertebrate datasets, mammals and ray-finned fish, and their corresponding USCO reference marker sets, actinopterygii_odb10 and mammalia_odb10. Our results show that although most USCO markers are single-copy, 85% and 91% respectively, empirical analyses using a large number of taxa are able to identify a non-negligible proportion of paralogs in the actinopterygii_odb10 and mammalia_odb10 BUSCO gene sets. Gene phylogenies show that several of these paralogs derive from ancient whole-genome duplication events, particularly the teleost-specific whole-genome duplication, and also more recent duplications such as those in salmonids and cyprinids. Paralogs do not reflect the organismal phylogeny but rather the history of gene duplications, so retaining paralogs in phylogenomic datasets can lead to inaccurate phylogenomic hypotheses. To test the effect of ignoring paralogy in BUSCO gene sets, we analyze the mammal and ray-finned fish datasets using concatenated maximum likelihood and summary coalescent methods by randomly choosing one paralog per locus if these are present. The resulting phylogenomic trees are overall consistent with the reference phylogenies, although small discrepancies were found. Phylogeny-guided paralogy removal also changes the sequences retained for many loci, but whether these changes alter the inferred species trees remains to be tested.

## Introduction

In recent years, advances in sequencing methods and computational technologies capable of analyzing vast amounts of data have enabled bioinformatics and related fields to progress by leaps and bounds. One such field is phylogenomics, which combines phylogeny and genomics to study the evolutionary relationships among organisms through genome analysis. By using DNA, RNA, or protein sequences, phylogenetic trees can reveal not only evolutionary relationships but also events such as gene duplications, horizontal transfers, or evolutionary convergence.

The lack of standardized marker sets for phylogenetic analysis leads researchers to use different gene sets, making results difficult to compare across studies. Certain marker genes are frequently employed due to their evolutionary characteristics, convenience, or historical reasons. Commonly used markers include genes encoding ribosomal RNA (*16S* and *18S rRNA*) and mitochondrial DNA (e.g., *cytochrome c oxidase subunit 1*). At the phylogenomic level, markers such as Ultraconserved Elements (UCEs) have been used (Faircloth et al., 2012). UCEs are, highly conserved DNA sequences across evolutionary lineages, or protein-coding genes derived from transcriptomes (phylotranscriptomic) or genome assemblies.

Recently, Universal Single-Copy Orthologs (USCOs) have been proposed as a set of nuclear marker genes that, due to their low duplication rates in genomes, are highly useful in phylogenomics. Their history begins in 2015, when they were first referenced (Simão et al., 2015) and defined as single-copy orthologs expected to be present in over 90% of selected species within a specific taxonomic group (e.g., mammals, actinopterygians). The latest release of BUSCO (*Benchmarking Universal Single-Copy Orthologs;* Tegenfeldt et al., 2025) includes 332 lineages for Bacteria, 29 lineages for Archaea, and 109 lineages for Eukaryota. The primary purpose of USCOs is to assess the quality of genome assemblies. A high number of complete USCOs, along with a low number of duplicates, fragments, or missing genes, indicates a high-quality assembly (Simão et al., 2015). In 2017, USCOs were proposed for phylogenomics (Waterhouse et al., 2017) using updated datasets and an improved version of the BUSCO software to identify orthologs that were subsequently used to infer phylogenies of birds, vertebrates, insects, and fungi. BUSCO identifies potential orthologs to reconstruct phylogenies with the reduced probability of introducing errors, e.g. paralogs (Simão et al., 2015; Waterhouse et al., 2017). Since the original paper, the usage of BUSCO have gradually increased from 43 citations in 2015 to 2720 citations in 2025 (query in Google Scholar was “BUSCO” + “Phylogeny”). USCOs were used to reconstruct phylogenies of Eumeninae wasps, combtooth blennies, monocots, and numerous other groups (Dai et al., 2023; Muñoz-Sánchez et al., 2026; Timilsena et al., 2022), with a recent large-scale assessment further demonstrating utility of BUSCO loci across diverse plant, fungal, and animal lineages (Alam et al., 2025). The choice of conserved markers can also affect which parts of the genome are represented. Foley and Murphy (2026) found that UCE and BUSCO loci are unevenly distributed across mammalian genomes and may fail to recover the species tree in clades with extensive introgression, underscoring the importance of marker choice in phylogenomic analyses.

In phylogenomics, accurate inference of evolutionary relationships among species relies on the correct identification of orthologous sequences. Orthologs are homologous genes that derived through a speciation event, while paralogs are homologous genes that arose through gene duplication (Figure 1). Traditionally, to find orthologs, the standard methods use: sequence similarity, gene trees, or a combination of both (Kapli et al., 2020; Altenhoff et al., 2016). OrthoFinder, for example, first clusters homologous genes into orthogroups and then analyses gene and species trees to infer orthologs and duplication events (Emms et al., 2026). Because orthogroups may contain paralogs, belonging to the same orthogroup does not automatically mean that sequences are orthologs.Orthology inference remains imperfect, and methods differ in how they balance the recovery of true orthologs against the inclusion of incorrect sequences (Altenhoff et al., 2016).

**Figure 1.**
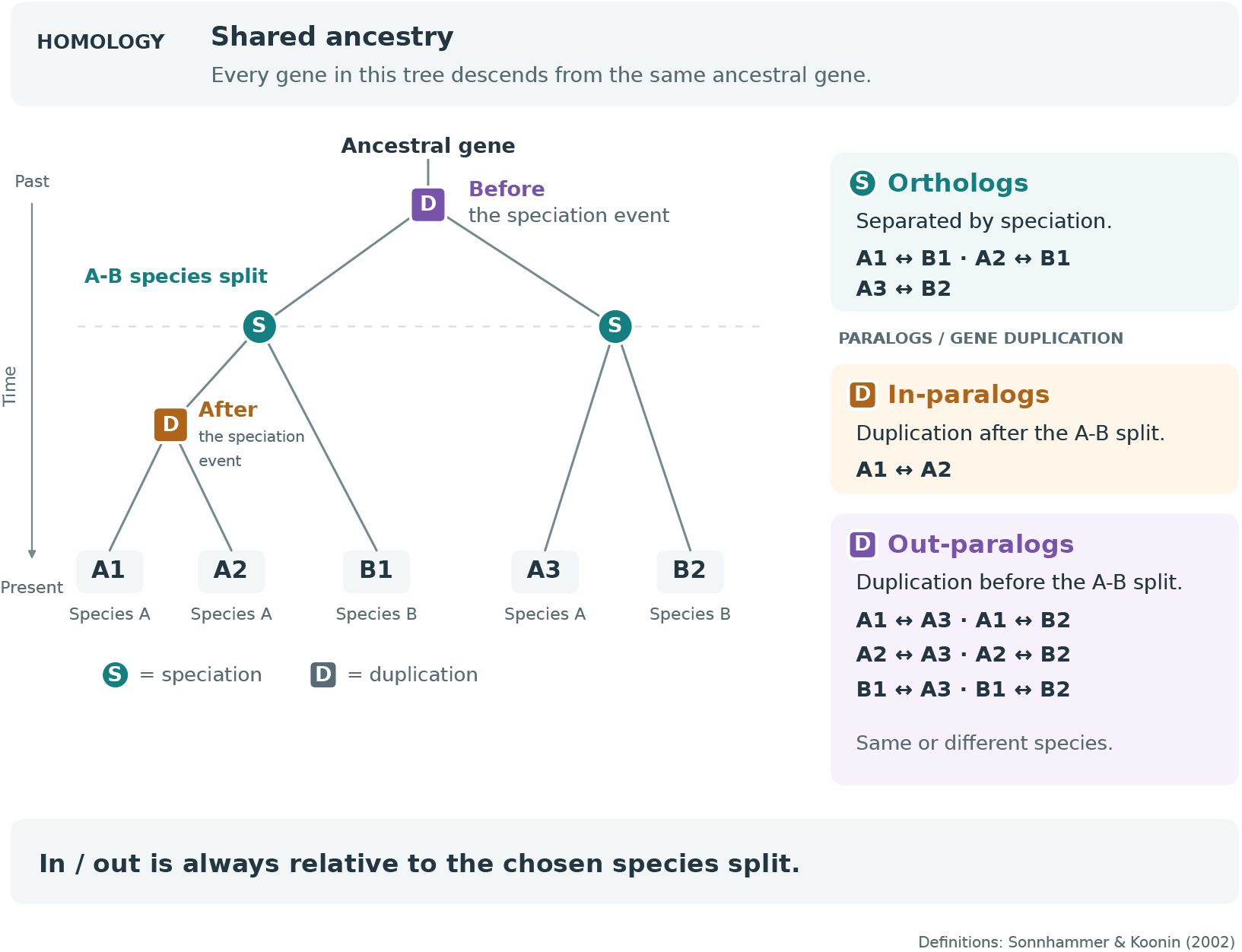
Gene relationships arising from speciation and duplication. All five genes are homologous. Terminology follows Sonnhammer and Koonin (2002).

Paralogs can be further classified as in-paralogs and out-paralogs according to the timing of the duplication relative to a reference speciation event (Sonnhammer & Koonin, 2002). In-paralogs arise from duplications occurring after the speciation event under consideration and out-paralogs result from duplications happened before this event. Thus, the distinction between in-paralogs and out-paralogs is inherently dependent on the speciation event used as a reference. In-paralogs generally do not violate the orthology assumption because copies arising after the relevant speciation event are co-orthologous to sequences from the other sampled species. Thus, retaining one representative in-paralog preserves the relevant orthology relationships, although the choice of copy may influence phylogenetic inference. In contrast, randomly selecting an out-paralog can violate the orthology assumption, as they originate from duplication preceding speciation event and should therefore be avoided. (Koonin, 2005).

On one hand, the information contained in sequence alignments is often insufficient to reliably differentiate orthologs from paralogs. On the other hand, when working with large-scale genomic datasets, we typically need to employ bioinformatic methods that may not faithfully detect all paralogy events. The retention of paralogs in a phylogenetic matrix can lead to incorrect resolution of specific nodes affected by higher duplication rates.

The inclusion of misidentified paralogs can alter phylogenetic relationships and produce a phylogenetic tree with high support of incorrect branches, although their impact appears to depend on taxon sampling (Struck, 2013). Later it was shown that even small proportion of paralogs could distort phylogeny (Shen et al., 2017; Irisarri, 2022).

Whole-genome duplication (WGD) is one source of such paralogs because it creates duplicate copies across many genes at once. Early vertebrate evolution included two rounds of WGD. Although most duplicated genes were subsequently lost during rediploidization, some copies were retained. These WGD-derived paralogs, known as ohnologs, preserve evidence of ancient duplication events (Simakov et al., 2020).

Beyond these two rounds of WGD, some vertebrate lineages have undergone additional round of WGD, such as the teleost-specific genome duplication (TGD) in ray-finned fishes. These WGD events not only drove vertebrate diversification but also left detectable patterns in chromosomal organization, revealing how gene duplication and reorganization shaped ancestral genomes (Simakov et al., 2020).

In the study by Parey et al. (2022), it was found that different species within teleosts had varying levels of ohnologs, suggesting that after the TGD, gene loss events were not uniform. This variability in the number of recent paralogs within each group can potentially bias phylogenies, altering inferences about relatedness and divergence between species. They showed that certain chromosomes underwent late re-diploidization events, meaning these ohnologs maintained very high similarity for millions of years, potentially causing problems in distinguishing between orthologs and paralogs. Ultimately, the TGD has increased the number of paralogs and complicates their evolutionary interpretation and the possibility of classifying them correctly (Parey et al., 2022).

Together, these duplication events raise a question for phylogenomics: how often BUSCO markers classified as single-copy across a reference set contain paralogs when sampled across a broader range of species? Our hypothesis is that the paralog prevalence correlates with the WGD events in actinopterygian lineage history and that retained paralogs might affect inferred phylogenetic relationships. We quantify the prevalence of paralogs, in-paralogs, and out-paralogs in the two USCO sets (mammal: Vargas & Zardoya, 2012; Foley et al., 2023; actinopterygian: Near & Thacker, 2024); identify actinopterygian species with elevated paralog counts; and compare the resulting phylogenies with reference phylogenies. We also examined whether lineages affected by additional genome duplication, including teleosts, show greater paralog prevalence. For both mammal and actinopterygian datasets we inferred maximum likelihood tree on concatenated tree and coalescent-based trees of USCOs to identify biases caused by paralog retention. Finally, we found substantial differences between USCO alignments produced by randomly retaining one sequence per species and those produced by phylogeny-guided paralog filtering.

## Materials and Methods

### Dataset assembly and gene-tree inference

Two phylogenomic datasets were assembled, one for mammals and one for ray-finned fish, using genome assemblies publicly available at the NCBI. To ensure a balanced taxonomic representation, we selected two species per taxonomic order following Vargas & Zardoya (2012) for mammals and Near & Thacker (2024) for ray-finned fish. This led to two datasets containing 45 and 78 species for mammals and ray-finned fish, respectively (Tables S1-S2). Mammalian species were selected to represent the orders described by Vargas and Zardoya (2012). For the reference phylogeny, mammal relationships followed Foley et al. (2023). For each genome, we identified USCOs using the BUSCO v5.7.1 in the ‘genome’ mode (Manni et al., 2021) and using the corresponding mammalia_odb10 and actinopterygii_odb10 lineage datasets. In mammal dataset for phylogenetic analysis, we retained *Ochotona princeps* (95.6% single-copy USCOs) instead of *Oryctolagus cuniculus* (91.0%) within Lagomorpha, and *Sorex araneus* (96.3%) and *Sorex fumeus* (96.6%) instead of Erinaceus europaeus (95.2%) within Eulipotyphla, leaving 43 mammalian species. All identified USCO genes (single-copy, duplicated, fragmented) were retained for subsequent analysis, comprising 9,224 and 3,640 USCOs for mammals and ray finned fish, respectively. Individual USCO genes were aligned using MAFFT v7.453we (Katoh et al., 2017). Alignment columns with >90% gaps were removed using ClipKIT v2.3.0 (Steenwyk JL. et al., 2020) in smart mode. Maximum-likelihood trees were inferred for each USCO from the filtered alignments using IQ-TREE v2.2.2.7 (Minh et al., 2020). The best-fitting substitution model was selected with ModelFinder (Kalyaanamoorthy et al., 2017) implemented in IQ-TREE according to the Bayesian Information Criterion (BIC). Branch support was assessed using 1,000 SH-aLRT replicates (Guindon et al., 2010).

### Species-tree reconstruction in a presence of paralogs

To test the effect of ignoring paralogy within USCO genes, we randomly selected one paralog when present in multiple sequence alignments using a custom Python script. This aims to mimic the situation when paralogy is ignored, and one of the paralogs is chosen based on non-phylogenetic criteria (e.g. the longest sequence). The corresponding reduced alignments (containing at most one sequence per species) were concatenated using AMAS v1.0 (Borowiec, 2016) into single matrices, for both mammal and ray-finned fish datasets. The concatenated matrices were analysed by maximum likelihood using IQ-TREE v2.2.2.7 (Nguyen et al., 2014) under BIC-selected best-fit models and 1000 pseudoreplicates of SH-aLRT. In addition, we inferred coalescent trees using ASTRAL-Pro 3 (Zhang & Mirarab, 2022) using individual USCO maximum likelihood trees as input, inferred as described above after the random selection of multi-copy sequences. To assess the possible effect of paralogy in BUSCO datasets, we compared the resulting concatenated and coalescent trees with the respective reference phylogenies of the original datasets (Foley et al., 2023 ; Near & Thacker, 2024).

### Phylogeny-guided paralogy filtering

We compared the USCO alignments used for species-tree inference, generated by randomly retaining one sequence per species, with alignments produced by a custom Python script for phylogeny-guided paralog filtering. When outgroup taxa was present in the gene, the gene trees were rooted to them: specifically to Ornithorhynchus and Tachyglossus for mammals and Erpetoichthys and Polypterus for fish. The midpoint-rooting was applied otherwise. Speciation and duplication events were inferred using species overlap algorithm (Huerta-Cepas et al. 2007) implemented in ETE4 (Huerta-Cepas et al. 2016). Duplication in which both branches contained sequences from the same single species were treated as in-paralogy, and the longest sequence was retained. All other duplications were treated as out-paralogy, for which the branch representing the largest number of species was retained and the alternative branch was removed from the alignment.

The example gene trees illustrating phylogeny-guided paralog removal are also shown.

## Results

### BUSCO recovery across genomes

We obtained a total of 9,224 USCOs for the mammal dataset containing a phylogenetically representative set of 45 species (the mammalia_odb10 lineage dataset contains 9,226 USCOs). Of these, 91,0% (min:62.5%, max: 97.9%, StDev: 6.1%) USCOs were, on average, single-copy. About 4.0% (min: 0.4%, max: 8.8%, StDev: 1.73%) were duplicated USCOs. On average, 1.6% (min: 0.7%, max: 16.4%, StDev: 2.4%) of USCOs were fragmented and 3.4% (min: 0.7%, max: 21.5%, StDev: 4.09%) were missing (Table 1). The actinopterygian dataset with 78 species representing all major orders was formed of 3,640 USCO loci (out of the 3,640 in the reference actinopterygii_odb10 lineage datasets). Of these, 84.6% (min: 10.9%, max: 99.3%, StDev: 17.45%) were on average single-copy, 5.1% (min: 0.1%, max: 68.0%, StDev: 13.29%) were duplicated, 4.4% (min: 0.4%, max: 23.7%, StDev: 6.27%) fragmented and 5.6% (min: 0.1%, max: 71.3%, StDev: 11.61%) missing.

**Table 1.** Statistics of USCOs after running the BUSCO program. The percentages represent the (rounded) average of the different types of USCOs identified by the program. The sum of “Complete” plus “Missing” plus “Fragmented” equals 100%, and within the “Complete” category are both “Single-copy” and “Duplicated” USCOs.

| Datasets | Mammals |  |  |  | Ray-finned fish |  |  |  |
| --- | --- | --- | --- | --- | --- | --- | --- | --- |
|  | mean | min | max | StDev | mean | min | max | StDev |
| **Total found USCOS** | 9224 | - | - | - | 3640 | - | - | - |
| **Completed** | 8765<br>(95,0%) | 5951<br>(64.5%) | 9088<br>(98.5%) | 542.0<br>1<br>(5.87%) | 3276 (90,0%) | 404<br>(11.9%) | 3622<br>(99.5%) | 635.1<br>(17.45%) |
| **Single-copy** | 8396<br>(91,0%) | 5766<br>(62.5%) | 9032<br>(97.9%) | 562.7<br>9<br>(6.1%) | 3079 (84,6%) | 397<br>(10.9%) | 3615<br>(99.3%) | 743.23<br>(20.42%) |
| **Duplicated** | 369 (4,0%) | 37<br>(0.4%) | 812<br>(8.8%) | 159.3<br>3<br>(1.73%) | 185 (5,1%) | 4<br>(0.1%) | 2475<br>(68.0%) | 483.88<br>(13.29%) |
| **Missing** | 314 (3,4%) | 65<br>(0.7%) | 1984<br>(21.5%) | 377.5<br>5<br>(4.09%) | 204 (5,6%) | 4<br>(0.1%) | 2595<br>(71.3%) | 422.72<br>(11.61%) |
| **Fragmented** | 148 (1,6%) | 65<br>(0.7%) | 1513<br>(16.4%) | 221.0<br>9<br>(2.4%) | 160 (4,4%) | 15<br>(0.4%) | 863<br>(23.70%) | 244.45<br>(6.72%) |

The higher proportion of fragmented (4.4% vs 1.6%) and missing (5.6 vs. 3.4%) USCOs in actinopterygians compared with mammals may reflect differences in genome assembly quality, which could reduce the recovery of complete USCO sequences.

### Distribution of paralogs

Classification of paralogs into in-paralogs and out-paralogs revealed substantial differences between actinopterygians and mammals (Table 2). Across the 3,640 USCOs in actinopterygians, 12,913 paralogs were identified, while for the 9,224 USCOs in mammals 7,880 paralogs were found (Table 2). These results gain even more relevance when we observe that only one third (3,089) of mammalian USCOs are affected by out-paralogs, while on the other hand, actinopterygians have 86% (3,144) of their USCOs affected by this process. The difference was also reflected in the overall number of paralogs per USCO, which was approximately fourfold higher in actinopterygians than in mammals (3.54 vs 0.85 paralogs per USCO, respectively).

**Table 2.** The total paralogs are the sum of both types of paralogs. The average number of paralogs (of any type per USCO) was calculated by dividing the total number of paralogs by the total number of USCOs.

| Datasets | Mammals | Actinopterygii |
| --- | --- | --- |
| Total found USCOs | 9224 | 3640 |
| Nºof genes with, at least, one *out-paralog* | 3089 (34%) | 3144 (86%) |
| Nºof genes with, at least, one *in-paralog* | 1455 (16%) | 2335 (68%) |
| Total paralogs | 7880 | 12913 |
| Total number of *out-paralogs* | 4745 (60%) | 6691 (52%) |
| Total number of *in-paralogs* | 3135 (40%) | 6222 (48%) |
| Number of USCOs with at least one paralog | 3790 (41%) | 3446 (95%) |
| Average of paralogs per USCO | 0,85 | 3,54 |
| Average of *out-paralogs* per USCO | 0,51 | 1,83 |
| Average of *in-* | 0,34 | 1,71 |
| ***paralogs per***<br>**USCO** |  |  |

We evaluated the number of paralogs in each individual species, particularly in actinopterygians. Table 3 shows the five species containing over 70% of the total actinopterygian paralogs.

**Table 3.**
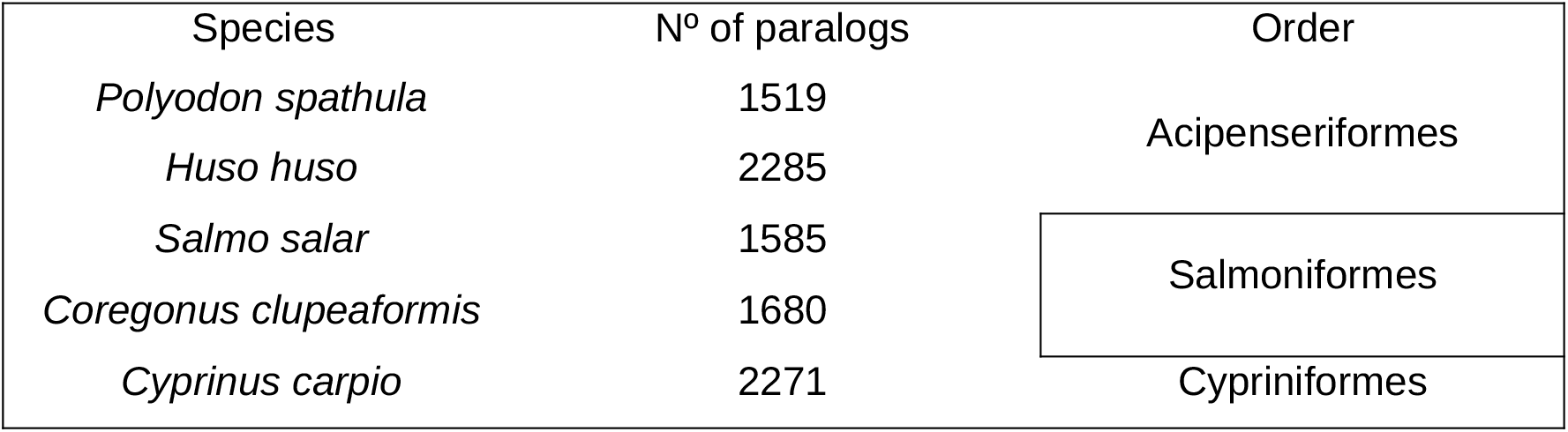
Actinopterygian species with high numbers of paralogs, resulting from specific duplication events.

| Species | Nº of paralogs | Order |
| --- | --- | --- |
| *Polyodon spathula* | 1519 | Acipenseriformes |
| *Huso huso* | 2285 |  |
| *Salmo salar* | 1585 | Salmoniformes |
| *Coregonus clupeaformis* | 1680 |  |
| *Cyprinus carpio* | 2271 | Cypriniformes |

### Species-tree inference

We have simulated the effect of ignoring the presence of paralogs in USCOs when performing phylogenomic analyses, randomly selecting one paralog per species in cases where they occur. We have inferred the phylogeny using two different methods: maximum likelihood and coalescence. In mammalian dataset, both the maximum likelihood and coalescence trees show a discrepancy in the order Artiodactyla, where *Sus scrofa* does not form the monophyletic group it should according to the reference phylogeny, with *Capra hircus*. The trees also differ from the reference in the placement of Perissodactyla: the reference groups it more closely with Carnivora and Pholidota, while our analysis place it closer to Cetartiodactyla. The mammalian reference phylogeny reduced to the set of taxa used in this study might be found in (Figure S1). The coalescence tree (Figure 2A), Perissodactyla shares a common ancestor with Artiodactyla and Cetartiodactyla, whereas in the reference phylogeny it does so with Pholidota and Carnivora. Most nodes in the coalescent tree have posterior probabilities greater than 0.95; the node associated with Dasyuromorphia has lower support (0.60).

**Figure 2.**
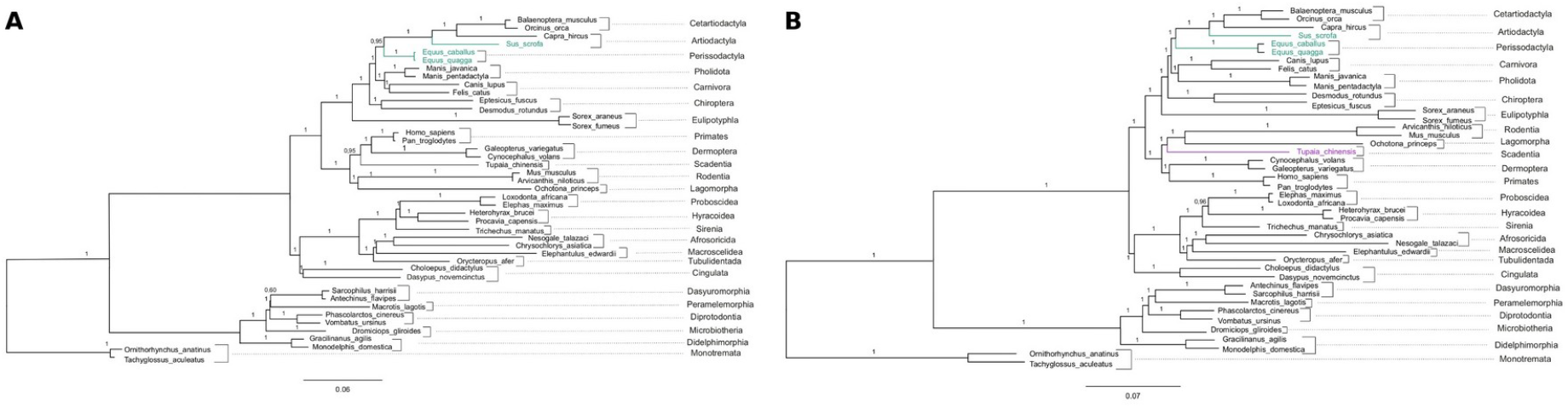
Mammalian phylogenies inferred using (A) the multispecies coalescent method implemented in ASTRAL-Pro3 and (B) maximum likelihood implemented in IQ-TREE2 using the concatenated alignment. Topological differences relative to the reference phylogeny are highlighted in green, while discrepancies between ASTRAL-Pro3 and IQ-TREE topologies are highlighted in purple. Taxonomic orders are indicated alongside the corresponding species.

The maximum likelihood tree (Figure 2B), Scandentia, represented by *Tupaia chinensis* (highlighted in purple), forms a monophyletic group with Lagomorpha and Rodentia. In both the coalescence-based and the reference phylogenies, it instead forms a monophyletic group with Primates and Dermoptera. All nodes have 100% aLRT support except the node separating Proboscidea, which has 96% support.

The actinopterygian phylogenies also differ from the reference tree (Figure 3). In both maximum likelihood and coalescence trees, the order Aulopiformes diverges earlier in the phylogeny than the order Ateleopodiformes (Figure 3), whereas the reverse order of branching is observed in the reference phylogeny. Syngnathiformes forms a monophyletic group with Gobiiformes in our trees but in the reference phylogeny, they form a monophyletic group with Scorpaeniformes. The final discrepancy between the likelihood and coalescence trees and the reference phylogeny concerns Acropomatiformes, which forms a monophyletic group with Centrarchiformes in our trees, whereas in the reference phylogeny, it is the sister group to Acanthuriformes. All posterior probabilities are above 0.85.

**Figure 3.**
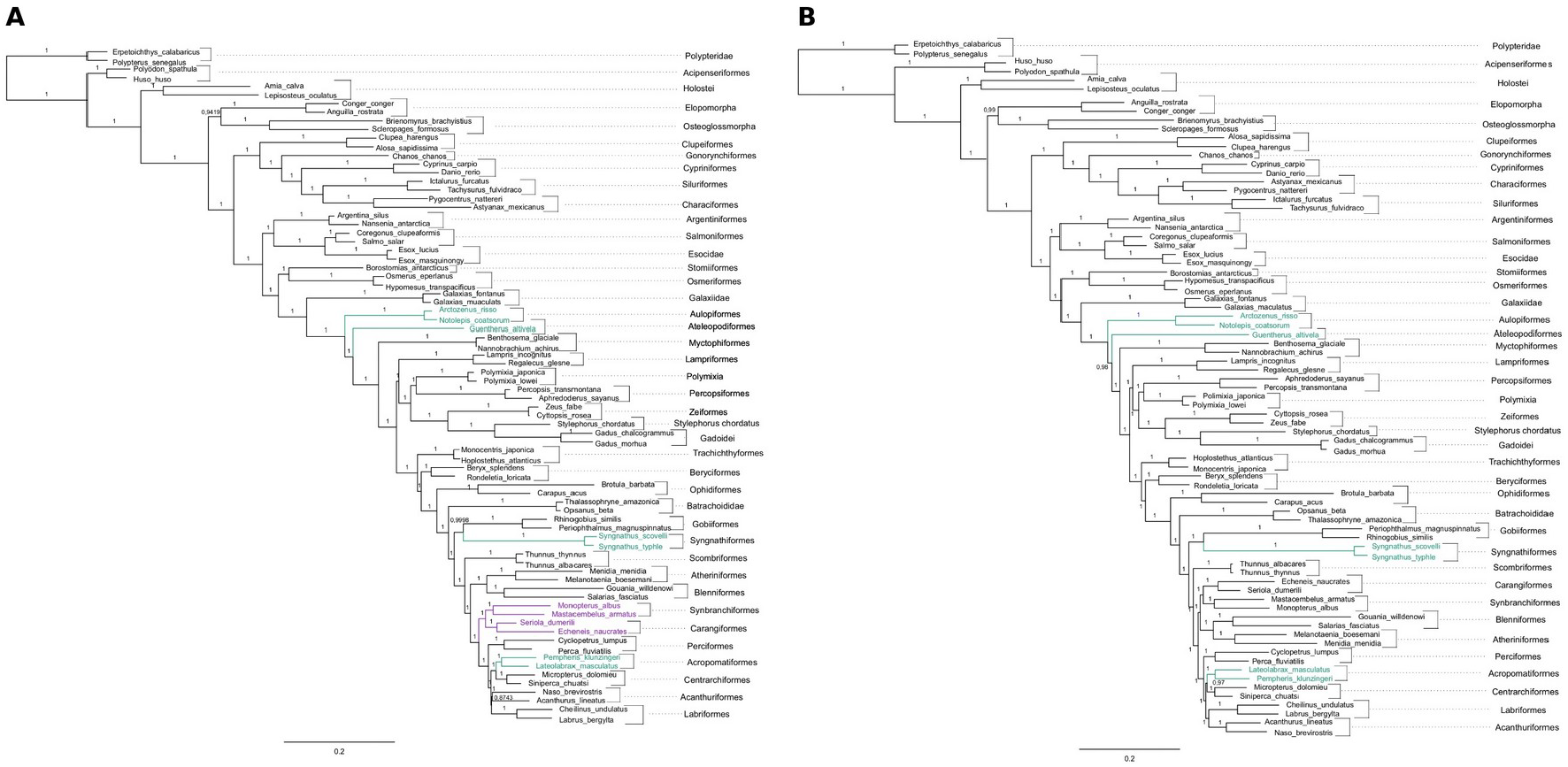
Actinopterygian phylogenies inferred using (A) the multispecies coalescent method implemented in ASTRAL-Pro3 and (B) maximum likelihood implemented in IQ-TREE2 using the concatenated alignment. Topological differences relative to the reference phylogeny are highlighted in green, while discrepancies between ASTRAL-Pro3 and IQ-TREE topologies are highlighted in purple. Taxonomic orders are indicated alongside the corresponding species.

The only topological difference between the maximum likelihood and coalescent trees is highlighted in purple in Figure 3. In the coalescent tree, the Carangiformes-Syngnathiformes clade does not form a monophyletic group with Blenniformes and Atheriniformes, whereas these taxa form a clade in both the maximum likelihood and reference trees. All aLRT values fall within the high-confidence range, being either 100% or very close to it.

### Phylogeny-guided paralog removal versus random sequence removal

For 6,675 (72.4%) of mammal genes and for 567 (15.6%) of fish genes, paralogy-filtering and random sequence removal produced identical alignments. Of these, 5,430 and 149 mammal and fish genes, respectively, required no removal (no paralogs found). In a further 1,245 mammals genes (13.5%) and 418 fish genes (11.5%), both methods retained the same sequences after filtering. Alignments differed in 2,545 mammal genes (27.6%) and 3,073 fish genes (84.4%). Mean retained-sequence F1 was 0.985 for mammals and 0.969 for fish. In 2,441 mammal and 3,049 fish genes, each method retained sequences absent from the other. In 104 mammalian and 24 fish genes, the random-removal set included all sequences by phylogenetic filtering and extra species (Figure 4).

**Figure 4.**
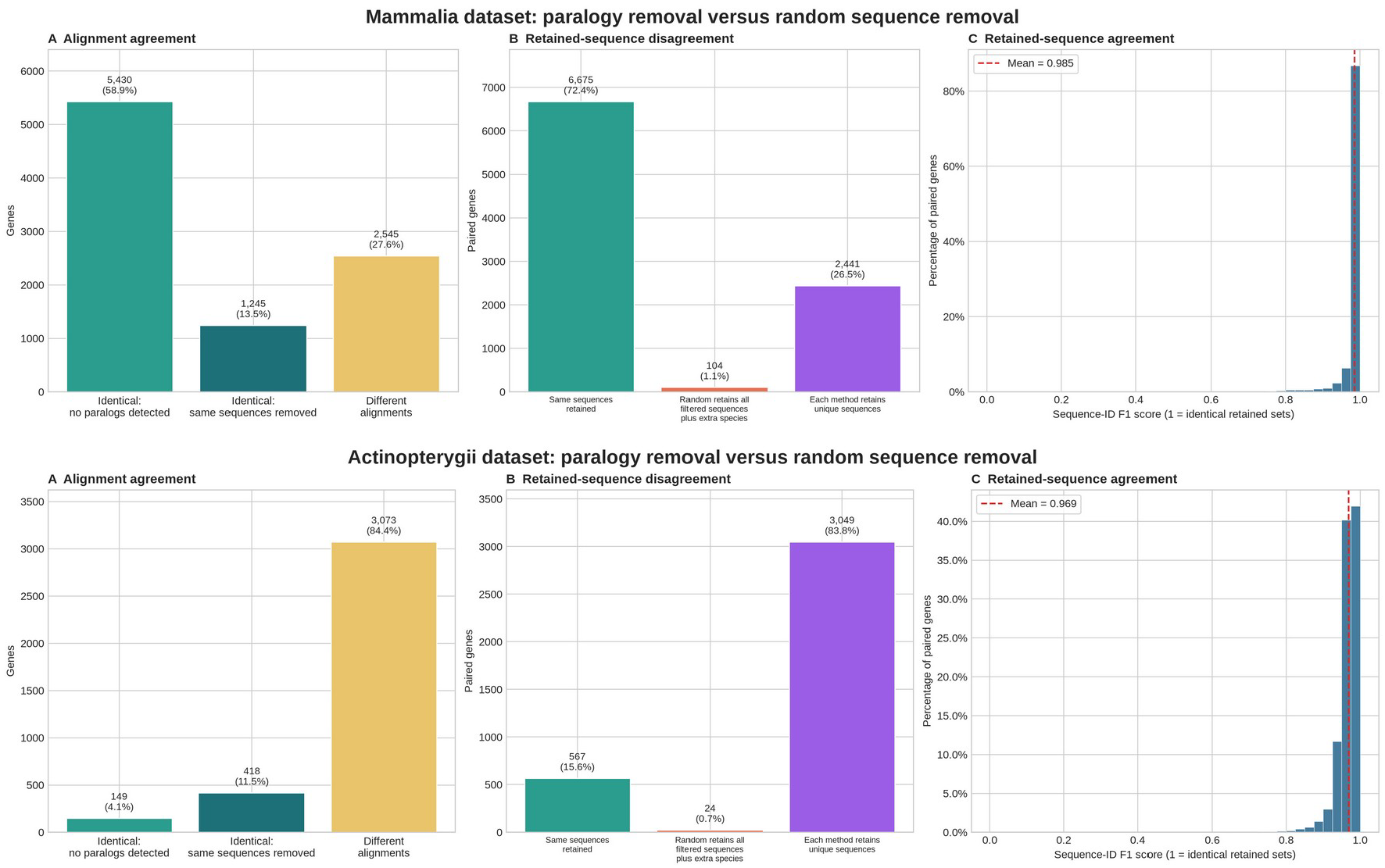
Comparison of phylogeny-based paralogy removal with random selection of one sequence per species in Mammalia (top) and Actinopterygii (bottom). (A) shows exact alignment agreement (B) shows whether the same or unique sequences were removed from alignments (c) shows per-gene F1 scores calculated from retained sequences.

Representative gene trees illustrate how phylogeny-guided filtering removed paralogs (Figure 5). In the mammalian gene tree, one in-paralog and one out-paralog were removed, reducing the alignment from 36 to 34 sequences. In the actinopterygian gene tree, three in-paralogs and four out-paralogs were removed, reducing it from 55 to 48 sequences. The latter also illustrates how filtering an out-paralog can remove an entire branch of a gene tree.

**Figure 5.**
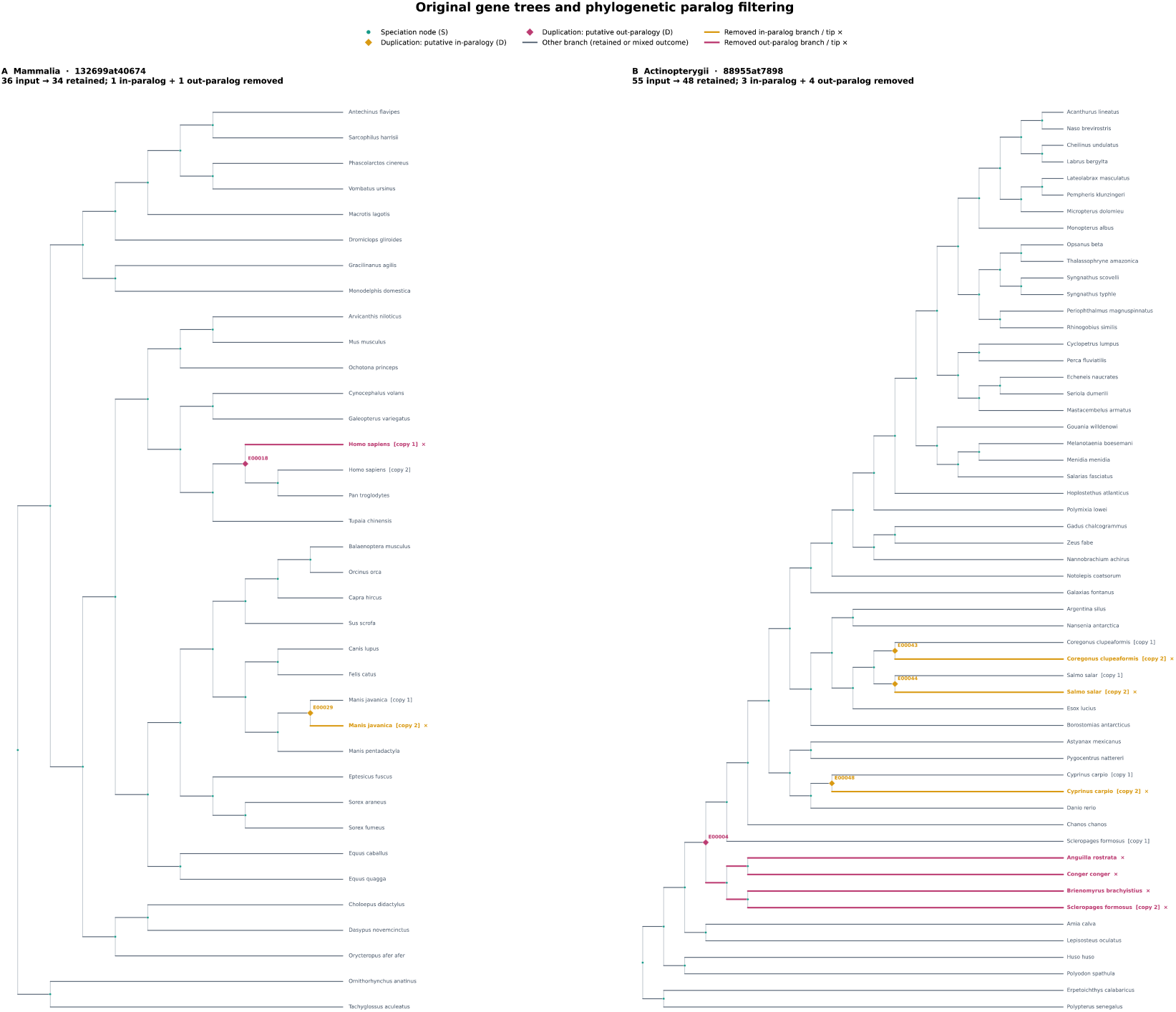
Example of paralog removal from mammal and actinopterygian gene trees.

## Discussion

We identified 9,224 USCOs in mammals and 3,640 in actinopterygians. This difference largely reflects the size of the lineage-specific BUSCO dataset: mammalia_odb10 contains 9,226 markers and actinopterygii_odb10 contains 3,640. This difference primarily stems from the evolutionary age of these groups: mammals represent a more recent group (283-296 Ma; Irisarri et al., 2017), making their genomes more similar and enabling the identification of a greater number of shared genes that meet USCO criteria. In contrast, actinopterygians have a much older evolutionary history (431-462Ma; Irisarri et al., 2017), leading to greater genomic divergence and fewer genes satisfying the strict requirements for USCO designation.

Among the 9,224 found mammalian USCOs, three (1155at40674, 151528at40674, 83074at40674) were represented in fewer than three of the 45 species and were therefore excluded from the phylogenomic analysis. The limited representation of these USCOs brings a question about their reliability as phylogenomic markers. Paralogy was more prevalent among actinopterygian USCOs than among mammalian USCOs: 95% versus 41% had at least one inferred paralogy event, and 86% versus 34% had at least one out-paralog. One possible explanation for this disparity might be in their distinct histories of gene duplication events.

Actinopterygians have undergone multiple ancestral genome duplication events throughout their evolutionary history, most notably the Teleost Genome Duplication (TGD or 3R) that affected all teleosts – though this represents just one of several such events. Because that duplication preceded the divergence of many sampled teleost groups, retained copies could be classified as out-paralogs relative to those speciation events. Of the 12,913 total paralogs identified in actinopterygians, 9,340 (72.3%) are concentrated in just five particular species (Table 3). These species belong to orders that experienced additional genome duplications (4R), namely Salmoniformes and Acipenseriformes.

Salmoniformes underwent an additional lineage-specific whole-genome duplication following the teleost TGD, which likely contributes to their elevated paralog counts (Lien et al., 2016). Acipenseriformes, although not teleosts, also have a complex history of independent whole-genome duplication and polyploidization events (Du et al., 2020). Despite extensive rediploidization following the ancestral whole-genome duplication of Acipenseriformes, a substantial proportion of duplicated genes has been retained, potentially contributing to the elevated paralog counts observed in *Polyodon spathula* and *Huso huso* in our dataset.

Following ancient genome duplications (like the two vertebrate whole-genome duplications), genomes typically re-diploidize (Simakov et al., 2020). However, more recent duplication events often maintain polyploid states or retain higher numbers of ohnologs through partial re-diploidization (Du et al., 2020). This explains the persistent paralogy patterns observed in certain actinopterygian lineages.

The fifth species exhibiting an exceptionally high number of paralogs is *Cyprinus carpio*. In addition to the TGD, the *C. carpio* lineage went through a more recent fourth round of whole-genome duplication (4R), whereas other cyprinids included in our dataset, such as *Danio rerio*, did not undergo this additional duplication. This 4R event resulted from allopolyploidization has been estimated to have occurred 12.4 million years ago (Xu et al., 2019). Notably, *C. carpio* displays a higher paralog count than salmonids (which also experienced 3R and 4R). The difference may partly reflect the more recent timing of the *C. carpio* WGD and, consequently, the retention of the larger proportion of duplicated genes. The *C. carpio* genome underwent allopolyploidization, while the salmonids WGD considered to have resulted from autotetraploidization (Robertson et al., 2017; Xu et al., 2019) that may also contribute to the observed pattern.

In phylogenomics, multiple methods exist for establishing sequence homology and orthology, each with distinct advantages and limitations, yet none are error-free. Consequently, while paralog filtering is desirable, no consensus exists on optimal methodologies, leading to study-specific approaches. The necessity to analyze thousands of genes precludes manual alignment scrutiny, requiring bioinformatic tools for filtering. The lack of standardized quality criteria and limited bioinformatic resources often result in paralogs being overlooked in phylogenomic studies.

Although only orthologs can accurately represent species phylogeny, several studies suggest that limited paralog contamination may not necessarily distort inferred tree topologies, particularly for well-resolved evolutionary relationships. For instance, in vertebrate phylogeny reconstruction, one or two paralogs didn’t compromise overall resolution, though they did affect some contentious branches (Irisarri et al., 2017). However, for resolving complex evolutionary relationships, paralog presence can significantly hinder phylogenomic inference (Irisarri et al., 2022; Shen et al., 2017).

The phylogenomic analysis of USCOs seeks to investigate whether the previously observed presence of paralogs affects the phylogenies inferred from these data. Our analyses suggest that even with a high prevalence of paralogs among USCO genes and a relatively large number of total paralogs and out-paralogs (especially in actinopterygians), randomly selecting one paralog allows us to obtain trees topologically similar to the reference phylogenies. However, there are certain incongruences between our phylogenies and the reference ones, as well as between the maximum likelihood and coalescence trees. These differences are likely concentrated in branches representing more complex evolutionary relationships, e.g. placement of Acropomatiformes. The relevant branch has a posterior probability of 0.8743 in the coalescent tree and SH-aLRT support of 97% in the maximum likelihood tree.

There are more discrepancies in the mammal trees, despite the higher proportion of paralog-free USCOs. The mammalian phylogenies differ from the reference tree in the placement of the Artiodactyla taxa. Retained paralogs may contribute to this discordance, but other factors, including incomplete lineage sorting, errors in gene tree inference, could also play a role. In the maximum likelihood method only one split was different compared to the reference phylogeny of mammals, and the coalescence method showed one split difference in actinopterygii. Branch support was consistently high in both the ASTRAL and IQ-TREE phylogenies, with values exceeding 85% across most nodes, except for the placement of Dasyuromorphia (Figure 2).

As expected, using a large number of genes resulted in highly supported phylogenies, supporting the suitability of USCOs as markers for phylogenomic inference. Nevertheless, the occurrence of paralogs might explain the discrepancy between reconstructed and reference topologies, this effect yet should be estimated.

The coalescent tree shows fewer differences from the mammalian reference tree than the maximum likelihood tree, and the opposite true for actinopterygian. The mammalian reference phylogeny was inferred using coalescence method (Foley et al., 2023) that may partly explain the greater agreement between the ASTRAL-Pro tree and the reference tree. The actinopterygian reference (Near & Thacker, 2024) is a summary phylogeny from multiple studies, where the maximum likelihood method was used. Therefore, these results do not establish that either inference method is generally more suitable for USCOs.

The coalescent-based methods have been shown (Yan et al., 2022) some robustness to the presence of paralogs. Moreover, ASTRAL-Pro3 is expected to be more robust to the presence of paralogs because, unlike standard ASTRAL, it was specifically designed to infer species trees from multicopy gene families while accounting for gene duplication and loss (Zhang & Mirarab, 2022).

Another important aspect to highlight is which species were used to define the mammalian and actinopterygian USCOs. For the mammalia_odb10 dataset (used to establish mammalian USCOs), 24 species were employed, nine of which are present in our matrix. Regarding actinopterygians, the actinopterygii_odb10 consists of 26 species, seven of which appear in our data. As previously mentioned, our species were randomly selected aiming for two species per order. Having more species than those included in the reference set increases the probability of finding paralogs in our phylogenies. This could be one of the main reasons for the high number of recent paralogs, since adding species not present in the original matrix reveals clade-specific duplications. This effect may be particularly pronounced in our actinopterygian dataset, which includes many teleosts that share an ancient genome duplication, as well as lineages that underwent additional duplications. In mammals, although they generally lack specific genomic duplications, putative recent paralogy was particularly frequent in Vombatus ursinus, Elephantulus edwardii, Choloepus didactylus, and Macrotis lagotis.

BUSCO markers are highly useful for assessing genome assembly quality (Simão et al., 2015). Across the genomes examined here, approximately 90% of markers were recovered as complete, single-copy genes. Although six of 45 mammals and 25 of 78 of fish genomes fell below the strict threshold of USCO genes definition.

In 2017, USCOs were proposed for phylogenomic applications (Waterhouse et al., 2017). Our study confirms that USCO genes serve as valuable molecular markers for phylogenomics, yielding robust phylogenetic trees that closely match reference phylogenies. Furthermore, neither method (coalescence nor maximum likelihood) demonstrates superior performance over the other. We have shown that USCOs could serve for detecting WGD, those duplication lead to the redundancy of paralogs which in their turn can influence phylogenetic reconstruction even when affecting only a limited number of branches.

In addition, we have shown that alignments produced by randomly retaining one copy per species often differed from those produced by phylogeny-guided filtering, which can remove an entire gene-tree branch represented by fewer species. The differences between the retained alignments suggest that phylogeny-guided filtering could affect species-tree inference. Trees inferred from those filtered alignments are needed to determine whether it changes the relationships reported here.

## Conclusions

Actinopterygians USCOs indeed contain higher number of paralogs compared to the mammals, therefore more careful paralogy filtering might be required for them. Within the actinopterygian dataset, paralogs were particularly abundant in species from lineages that underwent additional whole-genome duplication. In general, for both datasets USCOs showed to be appropriate for a reconstruction of highly congruent phylogenetic trees even in presence of paralogs. Phylogeny-guided filtering changed sequences retained for many loci, particularly in actinopterygians. Species trees inferred from the filtered alignments are required to assess whether this filtering improves congruence with the reference phylogenies.

## Acknowledgements

Computations were carried out in the HPC cluster of the Leibniz Institute for the Analysis of Biodiversity Change (LIB) and we thank Alexander Donath and Sebastian Martin for support.

