## Supplementary figures and images for "Benchmarking Universal Single-Copy Orthologs for Phylogenomics: The Impact of Paralogy"

### Supplemental_Figure_1

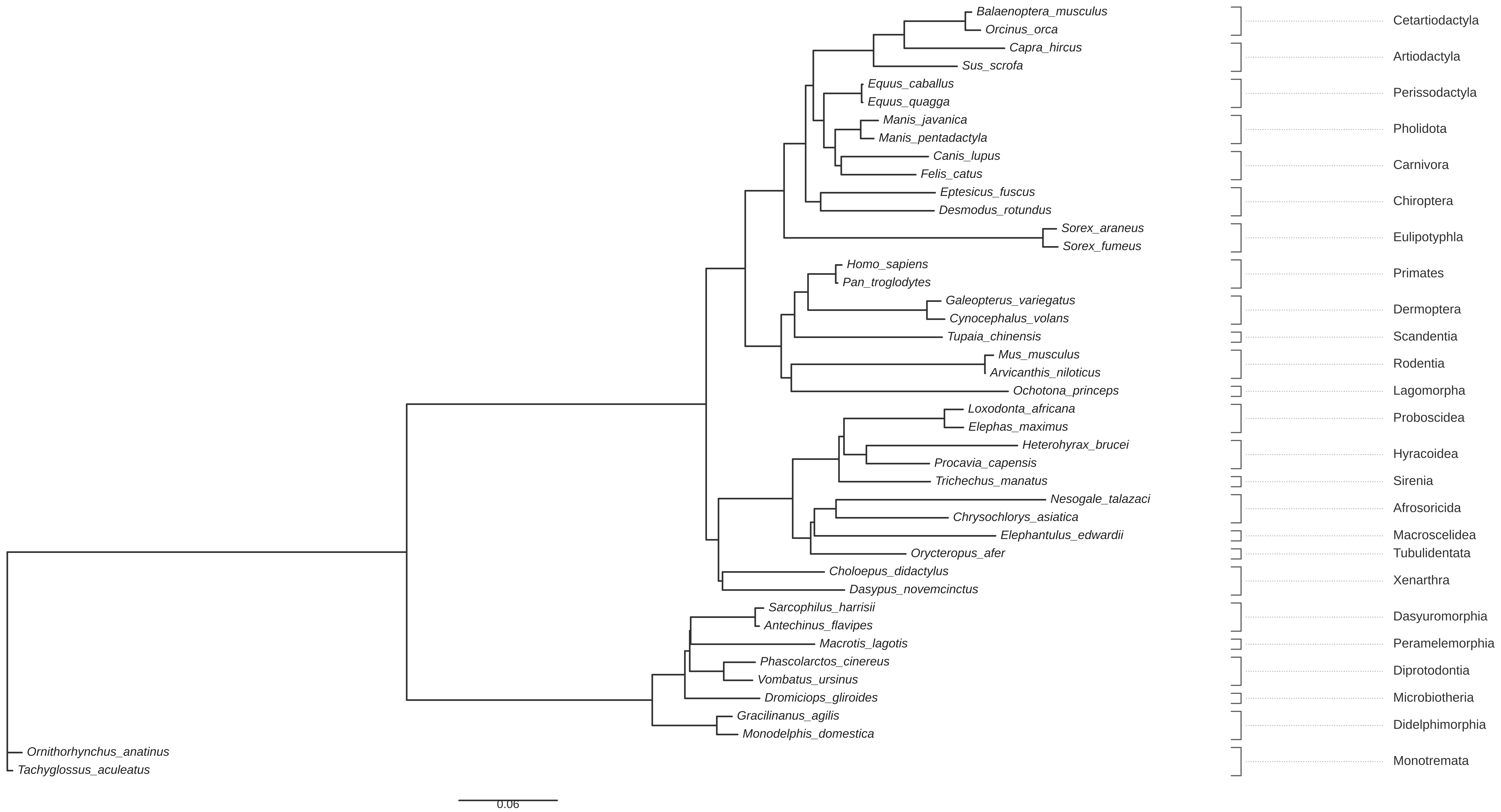
